# Detection of resistance in *Phytophthora infestans* to the carboxylic acid amide (CAA) fungicides using digital droplet PCR

**DOI:** 10.1101/2025.01.13.630889

**Authors:** Amanpreet Kaur, Ewen Mullins, Steven Kildea

## Abstract

**Background:** Potato late blight caused by *Phytophthora infestans* remains the greatest biotic threat to potato production globally. In northern Europe to prevent the disease and associated yield losses fungicides are heavily relied upon, with multiple applications required during the potato growing season. Unfortunately, such intensive fungicide usage has the potential to compromise control efficacy as it puts in place immense selective pressure for the emergence and spread of fungicide resistant strains of *P. infestans*. In recent years this scenario has been realised, with the emergence of strains resistant to the carboxylic acid amide (CAA) fungicides. As resistance to the CAA fungicides in *P. infestans* has been confirmed to result from changes in the pathogen’s cellulose synthase A3 gene (*CesA3*), specifically at amino acid position 1105, it opens the possibility to develop molecular tools to rapidly monitor populations of the pathogen for the resistance allele.

**Results:** Using the Stilla nacia 3-colour system a droplet digital PCR (ddPCR) was successfully developed to simultaneously detect the *P. infestans CesA3* gene irrespective of its CAA sensitivity status, and either the CAA wild-type sequence for position 1105 or that conferring the resistant allele G1105S. Using a ‘drop-off’ in ratio of positive droplets for either the wild-type or G1105S alleles relative to those positive for the general *P. infestans* CesA3 it was also possible to demonstrate that using the assay other potential alterations known to confer resistance that may occur at this position (e.g. G1105V) can be detected. The assay was validated using multiple sources of *P. infestans* DNA, including FTA cards.

**Conclusions:** The assay developed will allow for the accurate and sensitive detection of CAA resistance conferred by alterations at amino acid position 1105 in the CesA3 gene in *P. infestans*. The capacity to use the assay with multiple DNA sources and potential to adapt the ‘drop-off’ approach to different ddPCR platforms will ensure its applicability to the wider *P. infestans* community in monitoring the continued use of CAA-based fungicides for *P. infestans* management.

## Introduction

Potato remains a critical crop in the European cropping sector, and was estimated to be worth approximately €12 billion to the region in 2020(1). Even though it is over 175 years since the arrival of the oomycete *Phytophthora infestans*, the cause of potato late blight, the pathogen remains the biggest biotic threat to European potato production, costing the industry >€1 billion per annum(2). While reducing initial sources of *P. infestans* inoculum is vital for late blight control, it often falls short of providing the required season-long protection. In the absence of *P. infestans* resistant varieties, coupled with the rapid emergence of novel, virulent *P. infestans* strains, the intensive use of fungicides is a commonly used disease management strategy(3). Consequently, fungicidal control begins soon after crop emergence and continues until crop desiccation, sometimes requiring up to 20 applications per season in Europe(4). Thankfully the availability of various fungicide groups allows for targeted applications, which in turn is designed to maximize efficacy and longevity(5).

Amongst these fungicide groups are those belonging to the carboxylic acid amides (CAA) (FRAC Code no. 40) group, such as mandipropamid, benthiavalicarb and dimethomorph. By disrupting cellulose synthesis the CAAs inhibit cellulose production which the oomycetes rely upon to provide cell wall structure(6). Resistance to the CAAs was first reported in the grape pathogen *Plasmopara viticola* in the mid-1990s, following the introduction of dimethomorph(7). Subsequent studies by Gisi et al.(^7^) suggested that this resistance in *P. viticola* was recessive, with these findings later confirmed by Blum et al.(8). The authors found that resistance to the CAAs in *P. viticola* was conferred by a single mutation in the cellulose synthase 3 gene (*CesA3*) leading to the substitution of the amino acid glycine at position 1105 by serine (G1105S), and that the substitution needed to be in a homozygous state for the resistance phenotype to be expressed. CAA resistance has since also been detected in field populations of *Pseudosperonospora cubensis*(9), and in lab induced mutants for *P. infestans*(10), *P. melonis*(11), *P. capsici*(12) and *P. sojae*(13). In all instances resistance has been associated with amino acid substitutions in the respective pathogen’s *CesA3* gene, with substitutions of glycine at position 1105 most common. Similar to *P. viticola*, resistance in the lab induced *P. infestans* strain was recessive(10), whilst for *P. capsici* it appears to be semi-dominant(12).

Although *P. inf*estans can develop resistance to the CAA fungicides, the recessive nature of this resistance combined with the difficulties in generating resistance under lab conditions and the apparent absence of CAA resistance from field surveys, meant the risk of CAA resistance in *P. infestans* was deemed low(14, 15) (16). However, in 2019 initial reports of decreased efficacy of late blight control following the use of mandipropamid were first reported in Denmark, with subsequent field studies in 2020 and 2021 confirming this loss in efficacy(17). It has since been confirmed that this resulted from the presence of strains of *P. infestans* belonging to the lineage EU_43_A1 exhibiting high levels of resistance to mandipropamid(17). Strains of this lineage have since spread throughout Europe(18) (Hansen, 2023). As most late blight epidemics in Europe are driven by clonal populations of *P. infestans*, it is essential to identify potentially resistant strains quickly to allow strategies be implemented to minimise their selection within field populations. Traditional bioassays and molecular assays are routinely used to monitor fungicide resistance in oomycetes, as highlighted by Massie et al(19). Both assay types have advantages and disadvantages, often contrasting in speed, expense and specificity. For *P. infestans* the determination of fungicide sensitivity has until now almost exclusively been through the use of non-molecular based bioassays(17, 20, 21, 22). This has in part being due to the lack of information on the target sites of some of the key fungicides used for its control and/or the (potential) mechanisms of resistance that may be associated with these fungicides. The emergence of CAA resistance in *P. infestans* and confirmation that this resistance is associated with G1105S in the *PiCesA3* gene means it should be possible to screen *P. infestans* populations for CAA resistance through molecular techniques.

Such methods including PCR-RFLP, qPCR and pyrosequencing have been used to determine CAA resistance in *P. viticola* (23–25). Yet, PCR-RFLP and pyrosequencing are unfortunately limited in their detection capabilities, whilst qPCR is reliant on the production of a standard curve and as such it can be subject to the quality of the test sample. To overcome these limitations the concept of digital PCR (dPCR) was developed by Vogelstein and Kinzler(26). By separating a test sample into hundreds, thousands or even tens of thousands of separate nano-samples, with each individually undergoing PCR and end-point analysis, it is possible via Poisson’s distribution to calculate the initial DNA copy number in the test sample. With the use of microfluidics it has been possible to increase the ease and number of partitions created, hence the concept of droplet digital PCR (ddPCR). Its applicability has been further increased with dual-labelled probes that allow for specific detection of target sequences. Ristaino et al. (27) have demonstrated that ddPCR is an extremely sensitive method for the detection of *P. infestans,* whilst Zulak et al. (28) and Battisini et al. (29) have demonstrated the precision of ddPCR in detecting resistance to the azole fungicides and the Quinone outside Inhibitors, respectively in *Blumeria graminis* f.sp. *hordei* and *Zymoseptoria tritici*.

Here we report the development of a highly sensitive digital droplet PCR based method to detect CAA resistance in *P. infestans* conferred by the PiCesA3 alteration G1105S. The assay has been designed to work across different sample types (mycelia and disease lesions stored on FTA cards) ensuring its applicability to current and future monitoring methods.

## Material and Methods

### *Phytophthora infestans* isolates, DNA and synthetic gene fragments

DNA was isolated from the CAA sensitive *P. infestans* isolate Kerry_1_2023 (called CesA3-WT hereafter) available in the Teagasc Crop Research Centre, Oak Park, Carlow. DNA from the CAA resistant *P. infestans* isolate Pi155 homozygous for G1105S (called CesA3-mutant hereafter) was kindly provided by Syngenta. Gene fragments (350bp) of the *PiCesA3* gene from nucleotides 3,151 – 3,501 were separately synthesised in the form of gBlocks^TM^ Gene Fragments by Integrated DNA Technologies (Leuven, Belgium) to provide standards of the wild-type sequence, the sequence corresponding to G1105S, and G1105V (used to represent an alternative potential amino acid change at position 1105 known to confer resistance to the CAAs in *P. infestans*(10)) (see Table S1). Following initial reactions each fragment was diluted to an optimised concentration of 0.00001 ng/µl.

### Assay design

To exploit the potential of the Stilla nacia 3-colour ddPCR Platform (Stilla Technologies, Villejuif, France) a multiplex PCR was designed with three probes, *P. infestans*-Gen labelled with Cy5 to allow the detection of *PiCesA3* irrespective of CAA sensitivity status, PiCesA3-WT labelled with FAM to detect CesA3-WT and PiCesA3-G1105S labelled with HEX to detect CesA3-mutant (see Figure 1), thus allowing for detection of additional potential alterations at position 1105 that may confer CAA resistance. The primers and probes were designed based on the wild-type PiCesA3 gene sequence (EF563995.1) using Primer3Web (https://primer3.ut.ee/), with further optimisation of the different probes including the addition of locked nucleic acids (LNA) using the OligoAnalyzer^TM^ Tool (https://www.idtdna.com/pages/tools/oligoanalyzer) (see Table 1).

**Figure 1.**
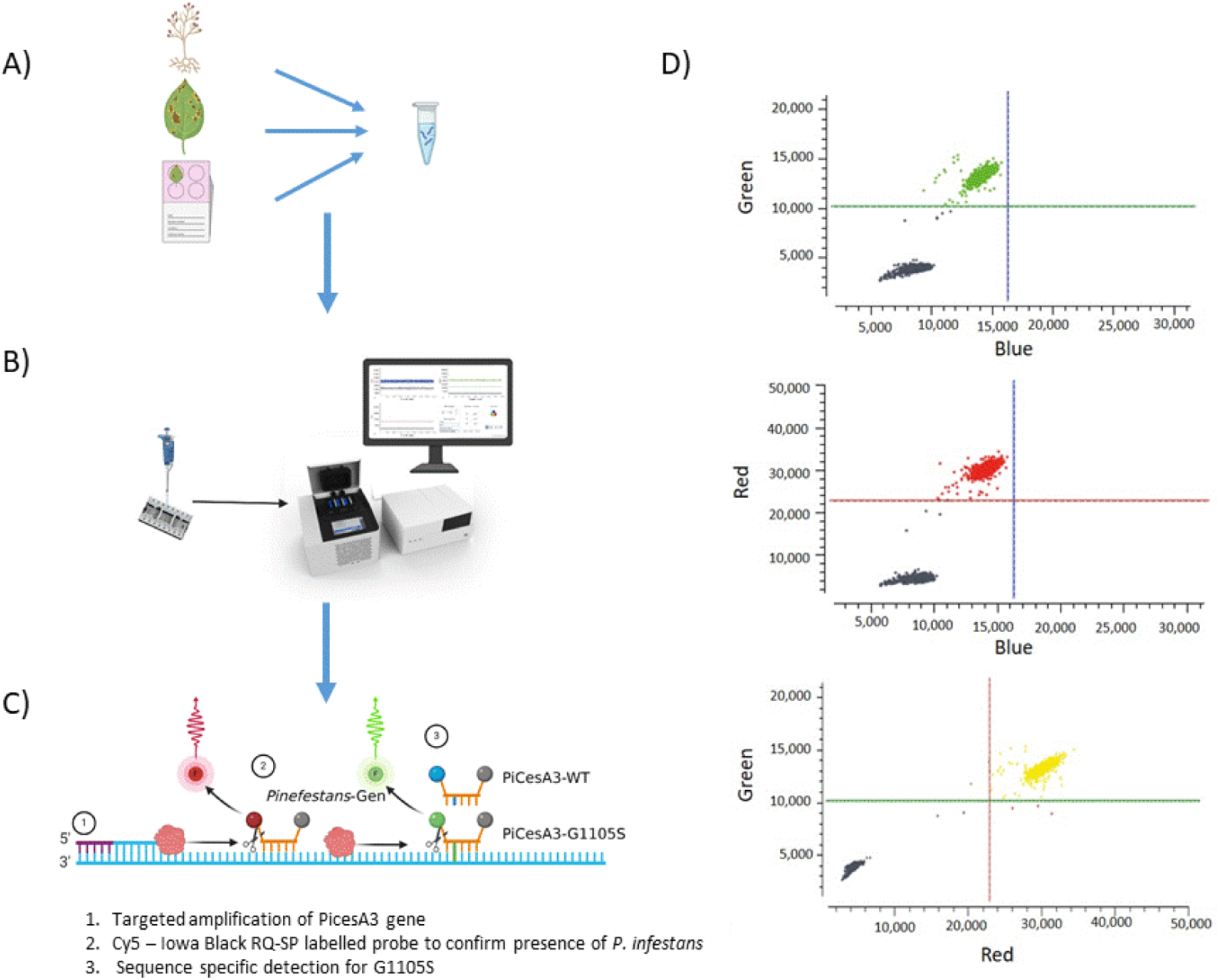
Overview of the ddPCR workflow to detect for CAA resistance in *Phytophthora infestans*. A) DNA is prepared directly from pure *P. infestans* cultures, or infected leaf tissue or infected leaf tissue stamped onto FTA cards following an initial nested PCR, B) DNA is loaded into a Stilla system and undergo sample partition and PCR, C) a combination of three dual-labelled probes are used to detect for changes that can confer CAA resistance, with G1105S specifically targeted, D) the presence of droplets positive for the *Pinfestans-*Gen probe, in combination positive/negative droplets for either PiCesA3-WT or PiCesA3-G1105S determines the sensitivity status of the specific droplet. In the 2D-fluorescence plots presented droplets are positive for both *Pinfestans-*Gen (Red) and PiCesA3-G1105S (Green) and negative for PiCesA3-G1105S (Blue), confirming the sample contains G1105S and is resistant to the CAA fungicides.

**Table 1:**
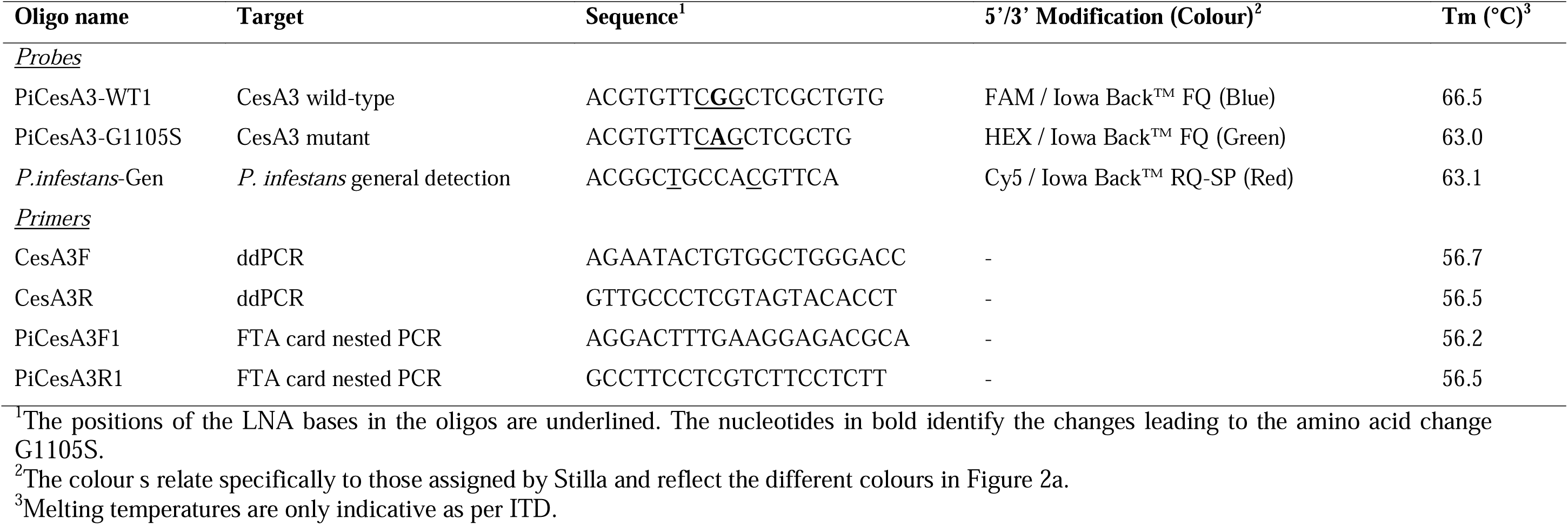
Primers, probes used for detection of substitutions in the *CesA3* gene associated with CAA resistance in *P. infestans*.

### Droplet digital PCR assay

ddPCR using the Stilla nacia^®^ system is divided into four specific elements. (1) Reaction mixtures are prepared containing naica^®^ multiplex PCR mix (1µl of Buffer A (5X) and 0.2µl of Buffer B (5X)), 0.50µl of each primer CesA3F/R (10 µM), 0.15 µM of each probe (10µM; CesA3-WT, CesA3-mutant and *P. infestans*-Gen), 1µl test DNA (DNA at 1 ng/µl, gBlocks as described previously at 0.00001 ng/µl or FTA PCR sample as described below), for assay optimisation and validation an additional 1µl potato DNA at a concentration10 ng/µl was added to the reactions to mimic the potential matrix of future test samples, and brought to a total reaction volume of 5µl with milliQ grade water. (2) The reaction mixtures were individually loaded into chambers of a Ruby Chip (Stilla Technologies) which was placed into the ddPCR machine (Geode), with a total of three Ruby chips (48 samples) possible per run. While in the thermocycler, sample partitioning takes place under pressure where samples are pushed through the channels in the chips for droplet generation. Each droplet corresponds to a single PCR reaction. The thermal cycling conditions were performed as follows: initial denaturation at 95 °C for 10 minutes, followed by 50 cycles of 95 °C for 15 seconds and 64°C for 1 minute (annealing temperature selected following optimisation as detailed below). 3) Once complete the chips were removed and scanned using a Prism3 multi-color fluorescence imager. 4) The acquired images were analysed using the Crystal Reader software (Stilla Technologies), where individual droplets were categorised as positive or negative for each of the three fluorescence channels, with absolute concentration or copies per µl (cp/µl) calculated.

Optimisation of the assay started by performing singleplex reactions for each probe and respective gene fragment. This progressed to multiplex reactions with all probes and single gene fragments, and finally singleplex and multiplex with mycelial DNA of known CAA resistance. The performance of the multiplex reaction as indicated by intensity of probe fluorescence and evidence of rain was tested at annealing temperatures 60°C, 62°C, 64°C and 66°C.

Limit of Blank (LOB) and Limit of Detection (LoD) were calculated for the detection of CesA3-WT and CesA3-mutant (α=5%) for each sample matrix separately in accordance to the protocol outline by Stilla Technologies (https://www.stillatechnologies.com/digital-pcr/statistical-tools/limit-detection/). Briefly, 30 replicate reactions negative for CesA3-WT or CesA3-mutant were run as described above. Copy numbers per µl were recorded per reaction, with reactions subsequently ranked in ascending order. To calculate the limit of detection (LoD), ten-fold serial dilutions of the samples (CesA3-WT and CesA3-mutant) 0.01 - 0.0000001 ng/µl were made and tested for positive signals to find a low level (LL) sample. Once the LL concentration was found, five independent LL samples (LL1, LL2, LL3, LL4, LL5) were prepared and six replicate reactions from each LL sample were completed and cp/µl determined. For both protocols the calculated cp/µl were uploaded on the Stilla Technologies LoB/LoD calculation software (https://www.stillatechnologies.com/digital-pcr/statistical-tools/limit-detection/).

To verify the linearity of the assay, gene blocks (CesA3-WT and CesA3-G1105S) were subjected to serial dilutions (with starting concentration of 1 ng/µl) and used as template in ddPCR. The actual concentrations (ng/µl) of the diluted samples were converted into cp/µl (Number of copies = (DNA concentration (ng/µl) x Avogadro’s number)/(gene length (bp) x conversion factor to ng x average weight of a base pair (Da)) and log value was calculated before plotting the graph.

### Assay specificity

To assess the specificity of the developed assay, pure cultures of eight *Phytopthora* species (*P. syringae, P. chlamydospora, P. cyptogea, P. cactorum, P. cambivora, P. pseudosyringae, P. lacustris, P. gonapodyides*) were obtained from the Plant Science Division of the Irish Department of Agriculture, Food and the Marine. Cultures were maintained on potato dextrose agar medium (Sigma-Aldrich, Germany) and 14 day old mycelium harvested, freeze dried and DNA extracted using a Qiagen DNeasy kit following manufacturer’s guidelines. The extracted DNA (1ng/µl) was used as a template in a ddPCR assay following above mentioned optimised conditions.

### ddPCR using FTA cards

FTA cards are critical for in-field sampling and monitoring of *P. infestans* in commercial potato crops across Europe (see https://agro.au.dk/fileadmin/euroblight/Protocols/Protocol_for_sampling_Phytophthora_infestans_DNA_using_FTA_cards_2020.pdf). To determine the feasibility of using the ddPCR assay to screen late blight lesions stored on FTA cards, 10 catalogued samples (from the 2023 season) previously genotyped as EU_6_A1, EU_8_A1, EU_13_A1, EU_36_A1, EU_37_A1, EU_43_A1 and characterised for the presence of G110S were used (Kaur et al., 2024). For this a nested PCR approach was taken, with an initial round of PCR to amplify a segment of the *PiCesA3* gene, used as template for the ddPCR reaction. Briefly a 2mm disc was cut from each sample and washed twice using FTA purification reagent (Whatman, GE Healthcare), before washing with TE buffer. The discs were dried at 37 °C for 20 minutes and used directly as a template for PCR amplification (7 cycles) using primer pair PiCesA3F1 (5’- AGGACTTTGAAGGAGACGCA-3’) and PiCesA3R1 (5’-GCCTTCCTCGTCTTCCTCTT- 3’) at Tm=60 °C. The PCR product was diluted to 1:10000 before being added as DNA template in ddPCR using the above optimised protocol.

To test the capacity of the ddPCR assay to detect multiple FTA samples in a single reaction, eight FTA samples, including a single sample confirmed as positive for G1105S, were combined post-*PiCesA3* template PCR and used as DNA template for the ddPCR. This pooling approach was then used to screen a total of 400 FTA stored *P. infestans* lesions taken from fungicide field trials conducted at Teagasc Oak Park in September 2023. Fungicide treatments included a range of fungicide modes of actions, including the CAA mandipropamid. Each FTA sample was processed separately and subjected to PCR amplification (as described above). 1µl from eight samples was subsequently combined together to make one pooled sample, which was screened using the ddPCR. This reduced the initial number of samples to be processed from 400 to 50.

## Results

### Optimization of the ddPCR assay

For all ddPCR reactions only samples that fractionated to >10,000 droplets were considered for the analysis. Initial reactions demonstrated that the different primer and probe combinations were able to amplify and identify their respective gene fragment targets when tested in singleplex and subsequently in multiplex (Figure 2a). There was some evidence of low levels of non-specific fluorescence for the CesA3-WT probe (FAM/Blue) and CesA3-G1105S (HEX/Green), however this did not interfere with the ability to distinguish between true positive and negative droplets. This separation was further increased by raising the annealing temperature to 64°C which was thereafter used as the optimised annealing temperatures for all reactions (Supplementary Figure 1).

**Figure 2.**
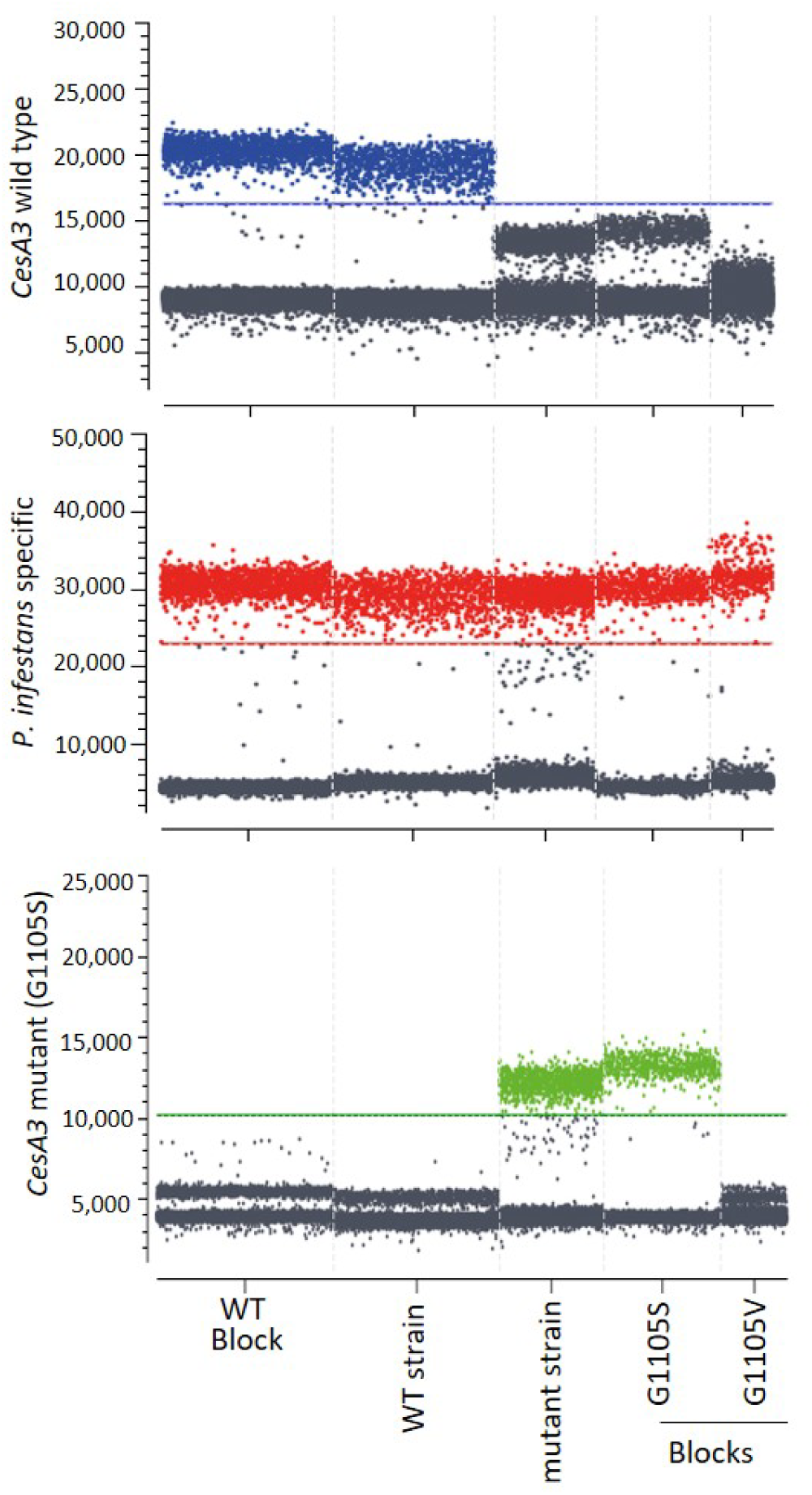
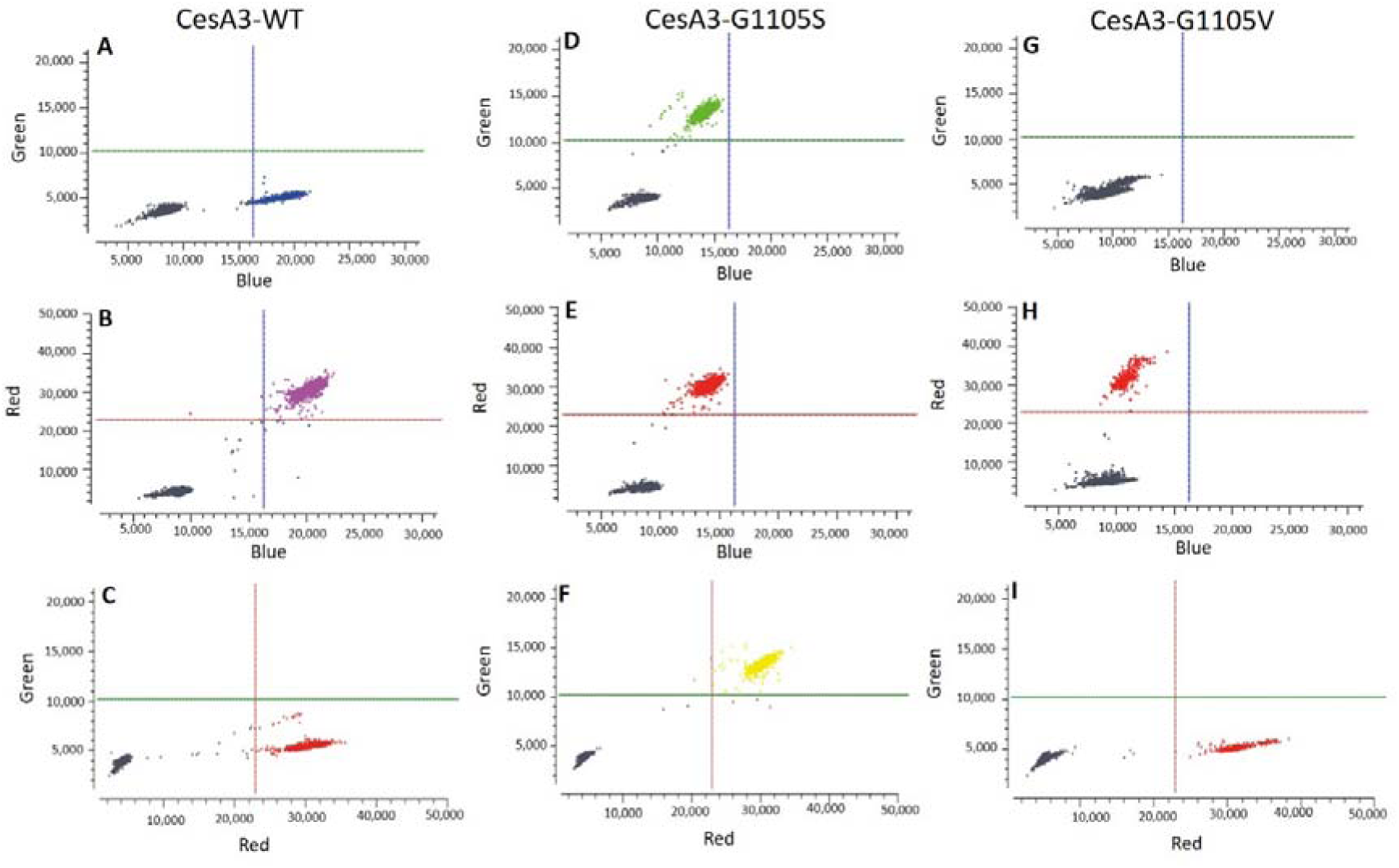
a) ddPCR fluorescence plots showing positive droplets for each of the three dyes (FAM/blue: PiCesA3-WT1, HEX/green: PiCesA3-G1105S, Cy5/red: *P.infestans*-Gen). Test samples include the synthesised 350 nucleotide gBlocks, the CAA wild type isolate Kerry_1_2023 and the CAA resistant isolate Pi155. b) 2D-fluorescence plots for detection of positive droplets in the Green-Blue, Red-Blue and Green-Red channels. Test samples were synthesised gBlocks containing the wild-type sequence for amino acid position 1105, the sequence conferring G1105S and G1105V. Combined they allow for the detection of A-C) wild-type *PiCesA3* which is positive in both the Red and Blue channels (Purple), D-F) the mutant PiCesA3-G1105S which is positive in the Red and Green channels (Yellow), G-I) the mutant PiCesA3-G1105V which is only positive in the Red channel.

As per the design of the assay (Figure 1), when applied in multiplex all reactions containing either *P. infestans* DNA (irrespective of CAA sensitivity) or spiked *PiCesA3* gene fragments had droplets positive in the Cy5/Red channel (*Pinfestans*-Gen probe). In reactions with gene fragments or *P. infestans* DNA with the *PiCesA3* wild-type sequence all droplets positive in the Cy5/Red channel were also positive in the FAM/Blue channel (highlighted as purple in Figure 2b), whilst gene fragments or *P. infestans* DNA with *PiCesA3* G1105S and positive in the Cy5/Red channel were also positive in the HEX/Green channel (highlighted as yellow in Figure 2b). For the G1105V fragment no further signals were detected in either the FAM/Blue or HEX/Green channels (Figure 2b), demonstrating the potential of the assay to detect other potential changes that may lead to CAA resistance.

Only four of the 30 reactions used to calculate the LoB for the detection of CesA3-WT showed any positive droplets, and amongst these all had <1 cp/µl. Therefore, the LoB for CesA3-WT was determined as 0.3 cp/µl. Using 0.0000001 ng/ul as the low level sample the LoD for CesA3-WT was calculated as 2.73 cp/µl (Supplementary Table1). For detection of the CesA3-mutant, positive droplets were detected in eight of the 30 reactions with <1 cp/µl in each of these; the LoB therefore was determined to be 0.51 cp/µl. Using 0.0000001ng/ul as the low level sample the LoD for CesA3-mutant was calculated as 2.20 cp/µl (Supplementary TableS1).

Further, the accuracy of the developed ddPCR methods was confirmed by determining the linearity of the assay (Supplementary Figure 2) which projects a linear relationship between observed and calculated copy number/µl for both CesA3-WT (R^2^ = 0.99) and CesA3-G1105S (R^2^ = 0.98) templates.

### Specificity of ddPCR to *P. infestans*

Nine species from the *Phytophthora* genus (*P. infestans*, *P. syringae, P. chlamydospora, P. cyptogea, P. cactorum, P. cambivora, P. pseudosyringae, P. lacustris, P. gonapodyides*) were used as template to evaluate specificity of the optimised ddPCR assay for *P. infestans*. Positive droplets above the LoD were only recorded for the *P. infestans* sample; irrespective of it being wild type or mutant (Figure 3).

**Figure 3.**
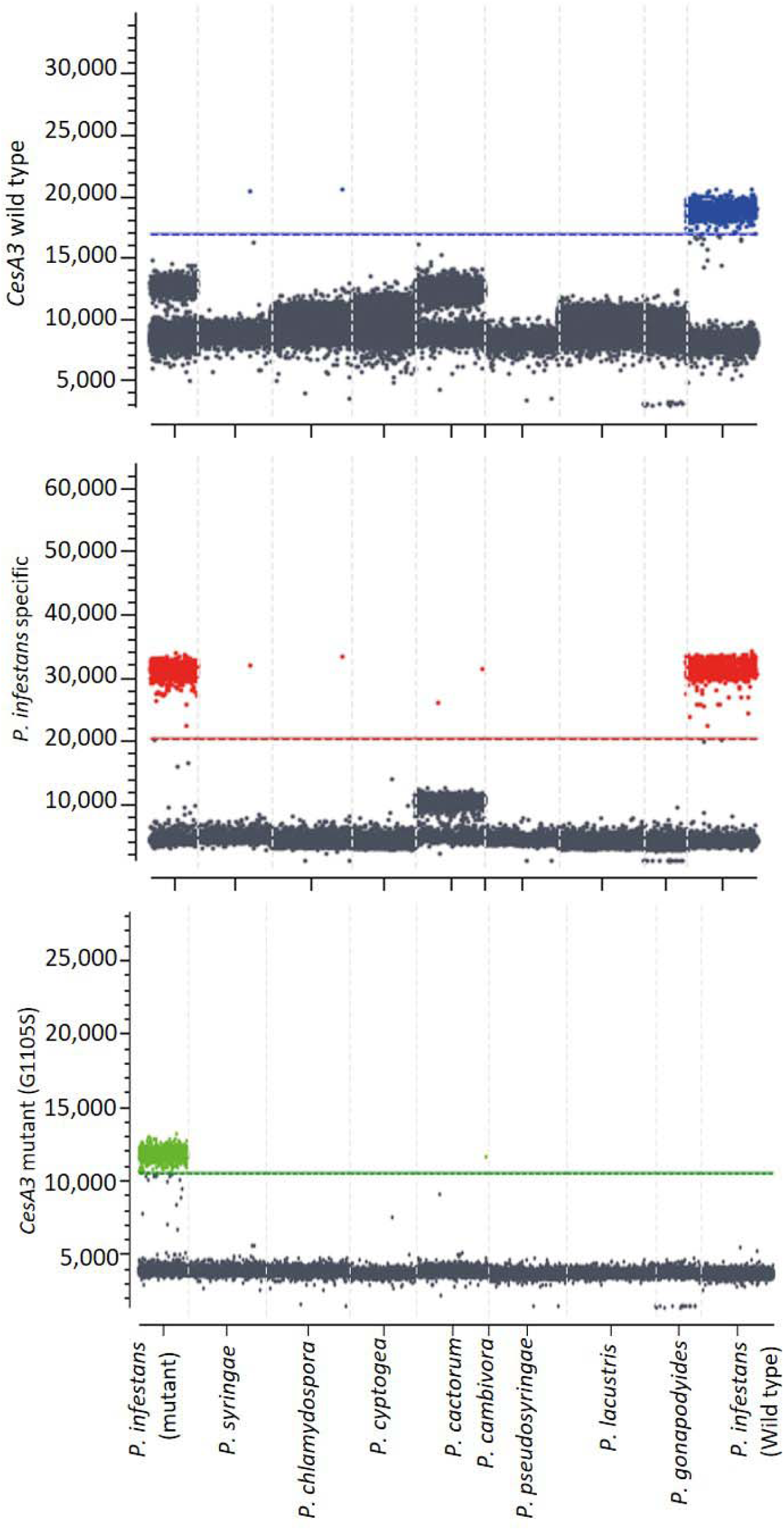
Testing the accuracy of ddPCR to detect *PiCesA3* gene in various *Phytophthora* species. *P. infestans* isolates Kerry_1_2023 and Pi155 carrying wild type and mutant *CesA3* gene respectivelywere used as controls

### Validity of ddPCR on FTA cards

Positive droplets for *P. infestans*-Gen (Red) were observed for each of the 10 FTA cards tested, indicating the positive detection of *P. infestans* on each card. For FTA samples belonging to the genotypes EU_6_A1, EU_8_A1, EU_13_A1, EU_36_A1, EU_37_A1 and previously confirmed as CAA sensitive, positive droplets were also observed for CesA3-WT (Blue). For the single sample belonging to genotype EU_43_A1, confirmed as CAA resistant, positive droplets were observed with the CesA3-Mut (Green) probe (Figure 4). When this sample was pooled with an additional seven known CAA sensitive samples it was still possible to detect its presence in the sample, demonstrating the capacity of the assay to be utilised to provide efficient initial CAA sensitivity screening of wider collections of samples. Amongst the 400 FTA cards from the 2023 growing season to which this pooling strategy was applied no CAA resistance detected, irrespective of fungicide treatment (Supplementary Figure 2).

**Figure 4.**
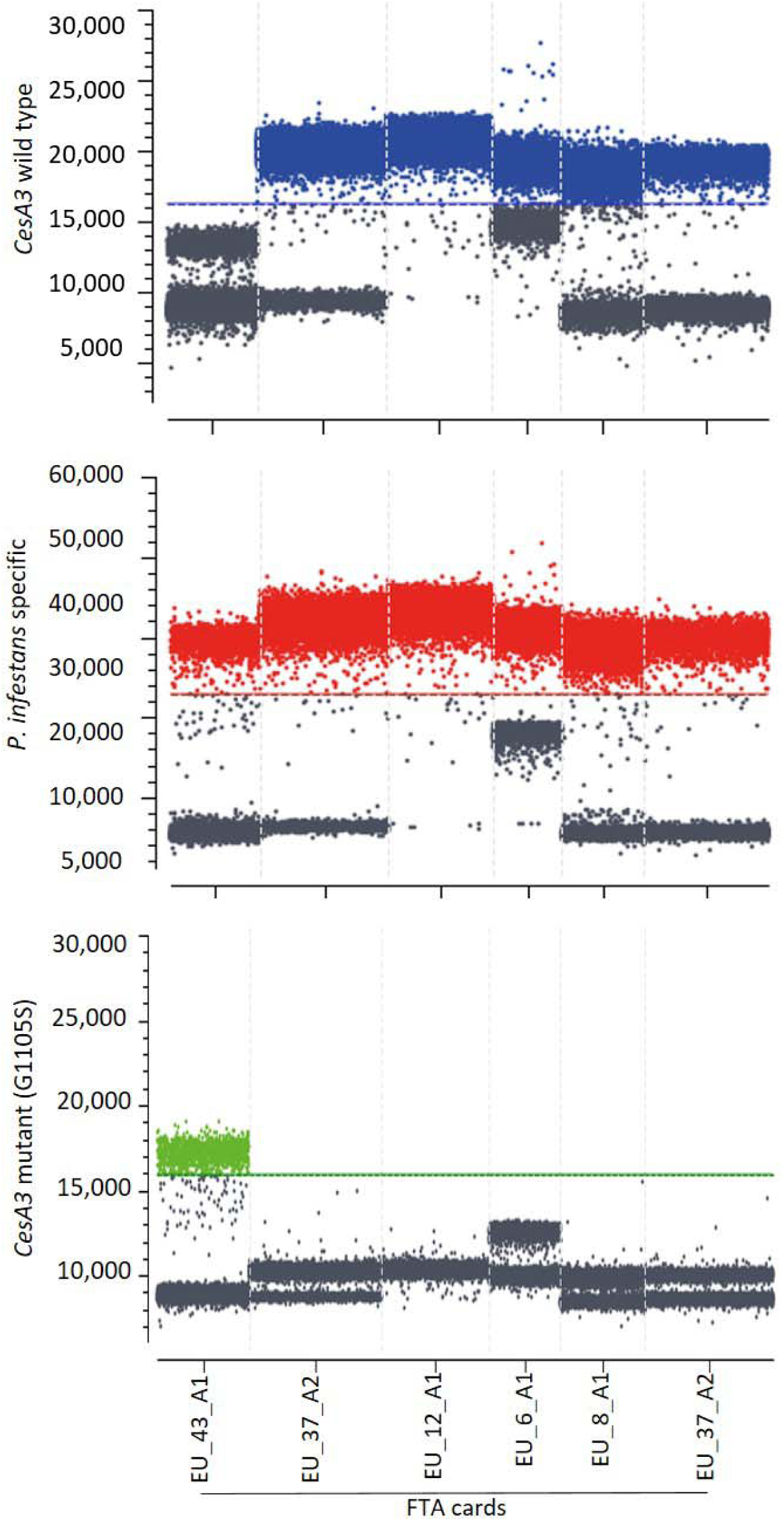
Detection of G1105S mutations in field samples stored as lesions pressed on FTA cards. Sample SK_24D (EU_43_A1 genotype) available at Teagasc OakPark as FTA card was used as a positive control.

## Discussion

The ability to monitor plant pests and diseases and to react accordingly through targeted control measures, such as fungicide applications, is a key pillar of integrated pest management (30). To date, for *P. infestans* this has primarily been based on genotyping using SSR profiles (31), with limited fungicide sensitivity screening (20). For the latter, the time and resources required to undertake such tests limit the capacity to use the information garnered within the season of collection. For the former, whilst it has the capacity to be used within season, it is currently limited to providing genotype lineage information. In some instances, such as fluazinam, a strong linkage has existed between *P. infestans* lineage and fungicide sensitivity phenotype (20) (21). In other cases, such as with CAA resistance the linkage with genotype is a bit more ambiguous. For example, to date all isolates of *P. infestans* that have been characterised as CAA resistant belong to the lineage EU_43_A1, however within this lineage strains sensitive to the CAA fungicides have also been identified (17) (32). As described by Abuley et al. (17) and Blum et al. (10) the fact that CAA resistance in *P. infestans* is conferred by alterations in the *Ces3A* gene means it should be possible to track these using molecular techniques, providing early and rapid detection of potential resistant strains. Such detection methods would undoubtedly aid in field decisions on fungicide selection

Using the Stilla Nacia 3-colour ddPCR platform we have demonstrated the ability to specifically detect the alteration G1105S in the *Ces3A* gene of *P. infestans* samples available as pure culture (mycelium) or FTA cards. By combining specificity for G1105S with the common detection for the *PiCesA3* gene in the same amplicon/droplet we demonstrate that it is possible using the assay to identify other potential amino acid alterations that may occur at position 1105, such as G1105V. Any drop-off in the number of droplets positive for either of the probes at this position relative to that for the common detection of *PiCesA3* indicates potential sequence variability in the region. The ability to perform this type of assay, referred to as a ‘drop-off’ assay builds on the fact ddPCR is based on the end-point quantification of droplets or partitions positive/negative for specific DNA sequences and has been successfully deployed in screening for mutational screening (33). To-date only G1105S has been detected in field samples of *P. infestans*, however as per Blum et al. (10) both G1105V/A have been confirmed under lab conditions to confer CAA resistance. Furthermore, in field populations of *P. viticola* and *P. cubensis* multiple alterations at position 1105 have been detected, demonstrating the capacity to detect other potential resistance conferring alterations at this position is important (34).

As most dPCR platforms are currently limited to two colours (35), by restricting the assay to the *P. infestans*-Gen probe (changing the attached probe to reflect system spectral range of the specified dPCR system if necessary) which detects *PiCesA3* irrespective of resistance status, and the CesA3-WT probe (FAM), which detects CAA wild-type *PiCesA3*, any differences in the number of droplets positive for the *P. infestans-*Gen probe and positive for both the *P. infestans*-Gen and CesA3-WT probes represent the frequency of resistance in a sample. In the absence of resistance conferring alterations at position 1105 the droplets positive for the *P. infestans*-Gen probe will also be positive for CesA3-WT.

To evaluate specificity, the optimized ddPCR assay was tested using nine *Phytophthora* species, confirming selective detection of *P. infestans* only. Whilst such specificity may not be required using potato field samples, whether using direct mycelia or samples stored on FTA cards, given the sensitivity of the assay it potentially could be used on spore samples as described by Arocha-Rosete et al. (36). Unfortunately, as CAA resistance conferred by alterations at position 1105 is recessive in *P. infestans* any such bulk sampling method will be limited to only confirming the presence of the resistance allele in the population and not the potential frequencies of resistance. Of course, if the frequency of the allele is >66% of the bulk sample it is possible to be confident that at least some proportion of the sample is CAA resistant.

The applicability of the assay for the early detection of G1105S in field samples was also demonstrated using a pooled FTA card approach. Using spiked samples positive amplification was only observed in samples containing either G1105S gene fragments or the *P. infestans* sample previously confirmed positive for G1105S and identified as belonging to the EU_43_A1 lineage (18). Using this pooled approach 400 x *P. infestans* infected potato leaves sampled from fungicide trials conducted at Teagasc Oak Park in 2024 were screened in a single ddPCR run, with all samples confirmed as CAA sensitive. As these samples were collected adjacent to that confirmed by Kaur et al. (18) as EU_43_A1, the failure to detect G1105S further confirms the belief that the initial detection of EU_43_A1 and G1105S was a secondary infection at the site, with a primary inoculum source elsewhere in the surrounding area.

The development of CAA resistance and the speed at which it has spread in northern and western Europe is a concern for the continued control of late blight, but equally for maintaining future fungicide anti-resistance strategies (17). A key component for both will be the ability to provide guidance on fungicide selection, with information on the fungicide sensitivity status of local *P. infestans* essential to achieving this. The development of the described ddPCR for the detection of G1105S and its application to field samples will aid these decisions. Whilst the CAAs are a key component of fungicide based late blight control, they are not the only fungicide mode of action to which *P. infestans* has developed resistance and additional assays (ddPCR or otherwise) to screen for resistance in the alternative MoA are urgently required.

## Acknowledgements

The authors would like to thank Dr. Lorenzo Borghi (Syngenta) for supplying the DNA of CAA resistant *P. infestans* strains, and Dr. Richard O’Hanlon (DAFM) for supplying the different *Phytophthora* species. This work was supported by Teagasc (project no. 0837) and the Research Leaders 2025 fellowship funded by European Union’s Horizon 2020 research and innovation programme under Marie Sklodowska-Curie grant agreement no. 754380.

---

**Supplementary Figure 1.**
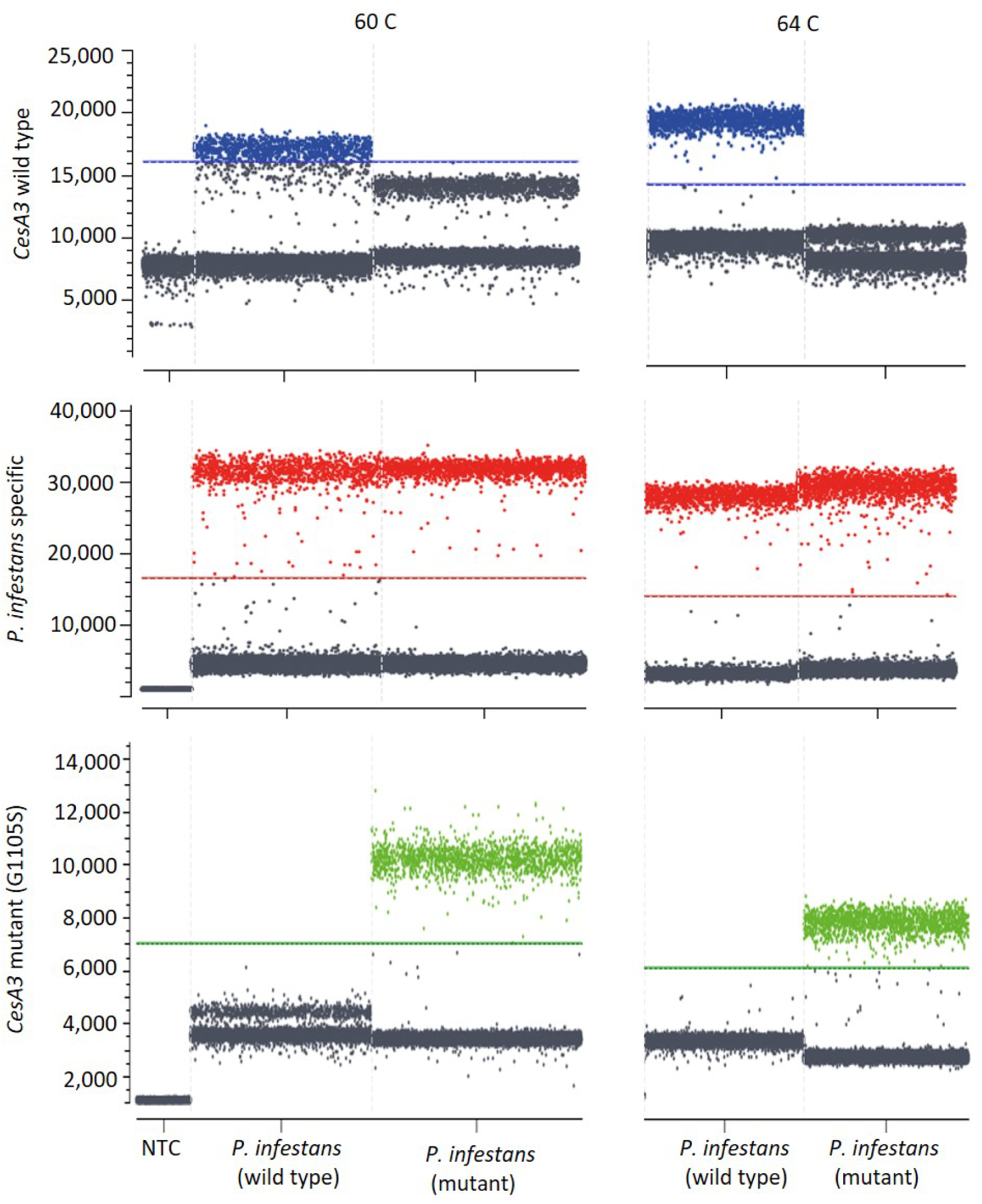
Temperature optimization of digital droplet PCR to differentiate between wild type and mutant *CesA* gene fragment in *P. infestans* species

**Supplementary Figure 2.**
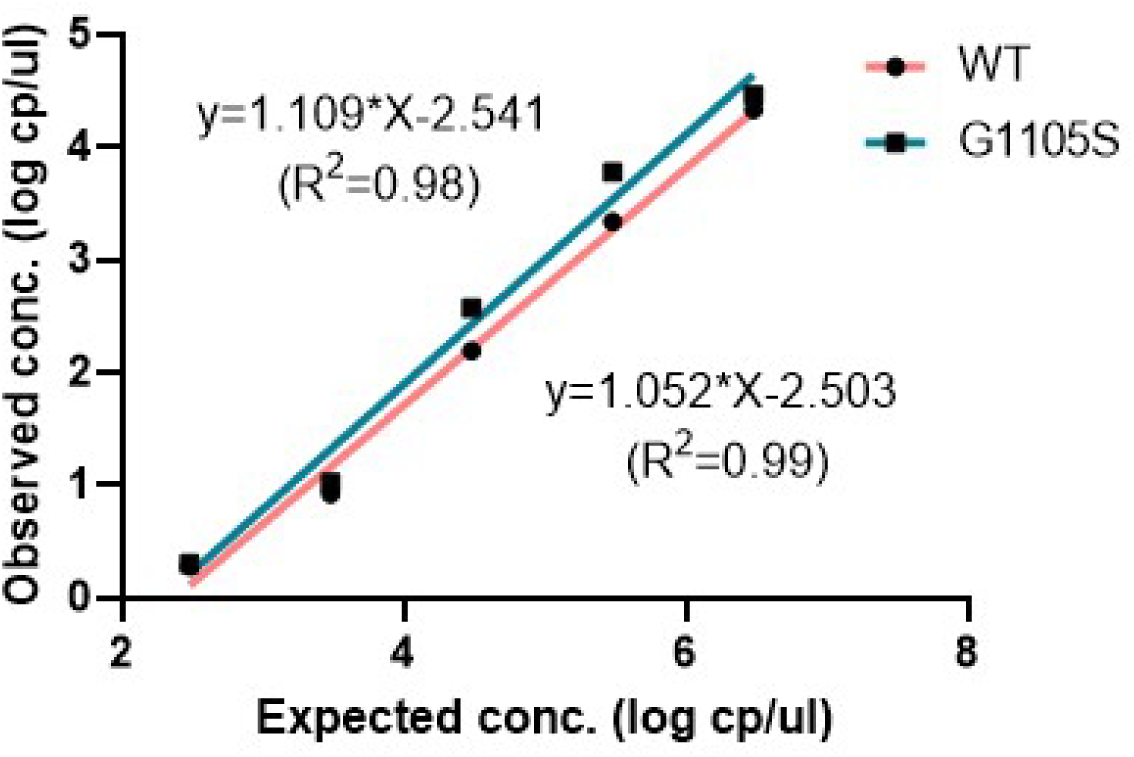
Linear range of the assay. Correlation between the concentrations observed in ddPCR and the expected concentrations.

**Supplementary Figure 3.**
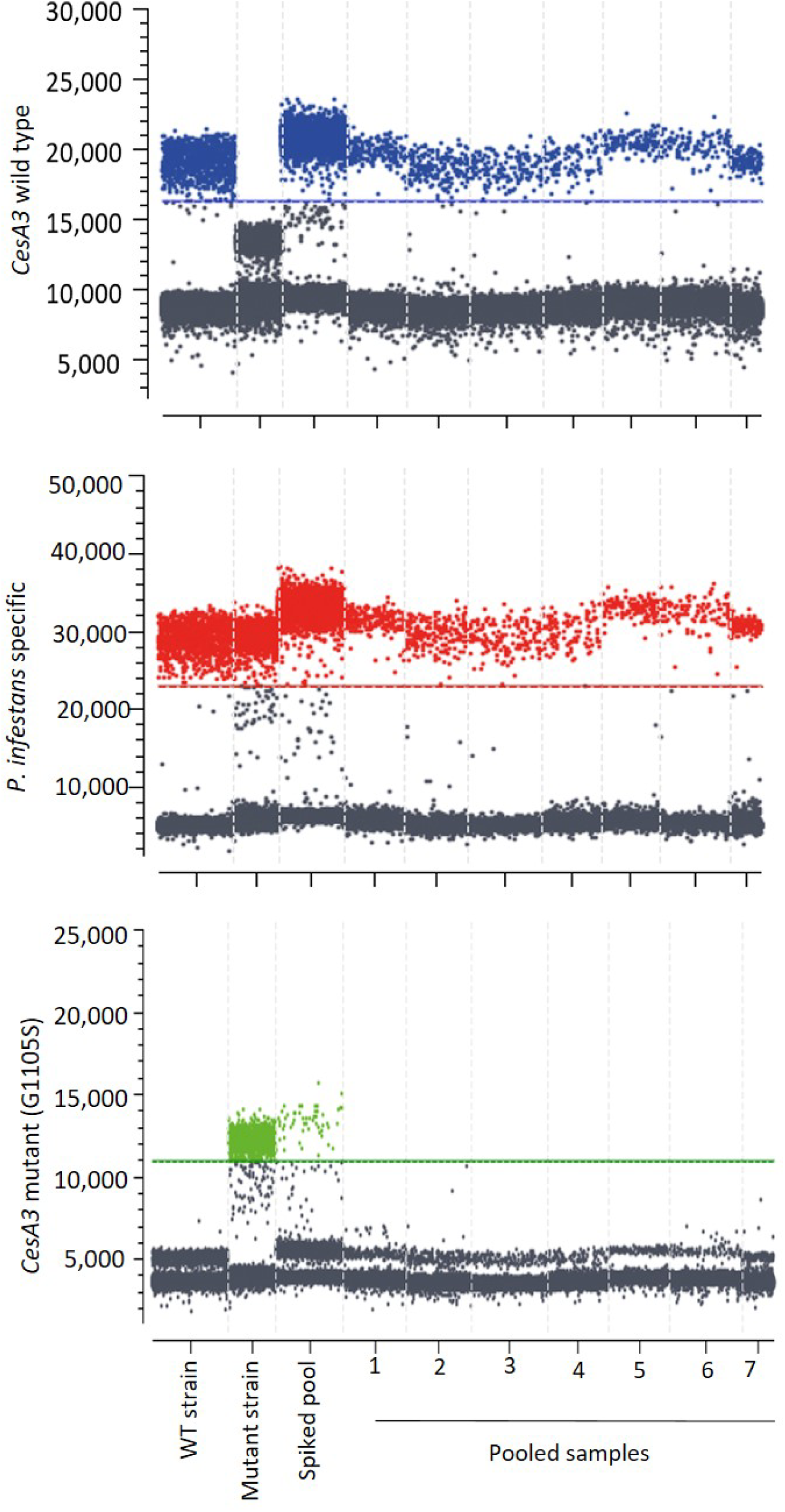
Ability of digital droplet PCR to detect the G1105S mutation in the pooled samples. Eight samples from nested PCR were pooled for testing. A sample known to contain G1105S mutation was included in a pool as a positive control

## Notes

### Competing Interest Statement

The authors have declared no competing interest.

